# Intrasegmental and propriospinal pre-phrenic interneuron circuitry in intact CNS and following cervical spinal cord injury

**DOI:** 10.64898/2026.09.08.749894

**Authors:** Pauline Michel-Flutot, Angela Harbeck, Lan Cheng, Khalil Rust, Megan A. Lyttle, Samantha J. Thomas, David A. Jaffe, Kallon Crowther, Gabriela Daszewska-Smith, George M. Smith, Shuxin Li, Angelo C. Lepore

**Affiliations:** Department of Neuroscience, Vickie and Jack Farber Institute for Neuroscience, Sidney Kimmel Medical College, Thomas Jefferson University, Philadelphia, PA 19107, USA; Department of Neuroscience, Shriners Hospitals for Pediatric Research Center, Temple University School of Medicine, 3500 North Broad Street, Philadelphia, PA 191405, USA

**Keywords:** cervical, diaphragm, interneuron, phrenic, propriospinal, respiratory, SCI

## Abstract

Cervical spinal cord injury (SCI) disrupts descending respiratory circuitry, resulting in debilitating and often persistent ventilatory deficits. Respiratory drive emerges within medulla from the rostral ventral respiratory group (rVRG), whose neurons project to C3-C6 phrenic motor neurons (PhMNs), which then innervate diaphragm, the primary muscle of inspiration. Though rVRG neurons make extensive monosynaptic connection with PhMNs, rVRG input to PhMNs can also be relayed through pre-phrenic interneurons (PP-INs) via polysynaptic pathways. However, the neuroanatomical connectivity between PP-INs and PhMNs remains incompletely understood. In both uninjured rats and the C2 hemisection (C2HS) model of cervical SCI, we performed tracing of PP-INs that were synaptically connected to PhMNs located in rostral (C3-C4) or caudal (C5-C6) portions of the phrenic nucleus by unilaterally injecting retrograde trans-synaptic tracer, pseudorabies (PRV), selectively into ventral or dorsal regions of hemi-diaphragm. We quantified numbers of PRV-labeled PP-INs individually at segments across cervical spinal cord, including separately in dorsal horn, intermediate gray, and ventral horn. We found that PP-INs are widespread throughout C1-C7 spinal cord in the uninjured condition. These PP-INs have a predominant intrasegmental connectivity pattern with PhMNs, though significant numbers of longer distance projecting propriospinal PP-INs also exist both rostral and caudal to the PhMN pool. Furthermore, while PP-INs connect both ipsilaterally and contralaterally with PhMNs, there is a strong bias to ipsilateral projection. C2HS induced major disconnection between PhMNs and ipsilesional propriospinal PP-INs located rostral to the SCI, but conversely induced limited plasticity in connectivity of intrasegmental PP-INs located within C3-C6 spinal cord. These findings greatly improve our knowledge about PP-IN circuitry and also provide important information to aid in developing approaches to target PP-IN plasticity for promoting spinal cord repair.

**HIGHLIGHTS:**

- Pre-phrenic interneurons are widespread throughout uninjured cervical spinal cord.
- PP-INs have a predominant intrasegmental connectivity pattern with PhMNs.
- Significant numbers of longer distance projecting propriospinal PP-INs also exist.
- There is a strong bias to ipsilateral connectivity of PP-INs with PhMNs.
- Cervical SCI induces major disconnection between propriospinal PP-INs and PhMNs.

## INTRODUCTION

Spinal cord injury (SCI) leads to persistent sensory, motor and autonomic deficits [1, 2], with cervical damage being the most prevalent in affected individuals. Cervical SCI disrupts descending respiratory pathways, resulting in long-lasting ventilatory dysfunction [3, 4]. To date, no treatment has been able to achieve complete spinal cord repair of respiratory circuitry, though some have shown substantial effect on functional recovery [5, 6]. In this context, deciphering how the cellular components of respiratory neural pathways interact in the normal condition and after cervical SCI is critical to support development of new therapeutics.

Respiratory drive emerges within medulla from the rostral ventral respiratory group (rVRG), whose neurons project to phrenic motor neurons (PhMNs) located in C3-C5/6 spinal cord. The axons of these PhMNs form the phrenic nerve that innervates diaphragm, the main inspiratory muscle [7, 8]. Though rVRG neurons are known to make extensive direct monosynaptic connection onto PhMNs [9], studies have shown that populations of interneurons within the spinal cord can also contribute to a relay between rVRG neurons and PhMNs, resulting in coexisting direct and indirect connections between bulbospinal drive and PhMNs. A large and highly diverse group of interneurons exists that is present at all spinal cord levels and at all gray matter laminae, is integrated into a wide range of spinal circuits, ranges from excitatory to inhibitory, and can project both intra- and intersegmentally [10]. Spinal interneurons involved in breathing that are integrated into phrenic motor circuitry and located in C3-C5/6 spinal cord were initially identified as “respiratory interneurons” [11, 12] and are now known as “pre-phrenic interneurons” (PP-INs) [13, 14].

Studies have begun to provide insight into these PP-INs. In addition to localization at spinal cord levels of the C3-C5/6 phrenic nucleus, PP-IN cell bodies also reside at high cervical levels C1-C2, and they can be found in dorsal, intermediate and ventral gray matter laminae across the various cervical segments [14]. PP-INs can project to and synaptically innervate respiratory motor neuron populations at-level (e.g., a C3 PP-IN synapsing with a C3 PhMN) and intersegmentally (i.e., a PP-IN cell body located at a different spinal segment than its post-synaptic target neuron), including with both ipsilateral and contralateral projections. Intersegmental PP-INs fall within the larger group of propriospinal INs, which are INs whose cell body resides in the spinal cord and projects its axon to a different spinal cord segment (either across just a single segment or for longer distances across multiple segments) [15]. Though these PP-INs can be identified experimentally using application of the trans-synaptic tracer, pseudorabies virus (PRV), to diaphragm [13], relatively little is known about their involvement in normal breathing and following SCI, as well as their potential recruitment when using therapeutics aiming at promoting spinal cord repair.

The C2 hemisection (C2HS) model in rodent is one of the most highly used tools to investigate cervical SCI and its impact on the respiratory system. C2HS disrupts descending bulbospinal input to ipsilesional PhMNs, resulting in diaphragm hemiplegia [13, 16–19]. To date, very few studies have investigated the response of PP-INs in cervical SCI paradigms such as C2HS [13, 20], highlighting the major gap in our knowledge regarding PP-IN biology in the pathogenesis of SCI.

In the present study, we extensively characterized both intrasegmental and intersegmental/propriospinal PP-IN circuitry within the cervical spinal cord of uninjured rats and following C2HS. Our findings provide extensive neuroanatomical insight to greatly improve our knowledge about PP-IN circuitry, including plasticity of their connectivity in response to cervical SCI.

## MATERIAL AND METHODS

### Animal studies

Female Sprague Dawley rats (ENVIGO, USA, n = 26, 250-300 g) were used for this study. All experimental procedures were approved by the Institutional Animal Care and Use Committee (IACUC) committee of Thomas Jefferson University and conducted in compliance with the National Institutes of Health (NIH) Guide for the Care and Use of Laboratory Animals and the ARRIVE (Animal Research: Reporting of In Vivo Experiments) guidelines. Rats were housed three animals per cage in a temperature, humidity, and light-controlled environment. Food and water were provided *ad libitum* on a 12-hour light/dark cycle. Only female rats were used in this study, as we have extensively optimized and comprehensively characterized the C2HS SCI model both functionally and histologically in female rats in our previous work [21, 22]. Animals were randomly assigned to group, and all surgical procedures and subsequent analyses were conducted in a blinded manner.

### C2 spinal cord lateral hemisection surgery

Twelve rats received a C2 spinal cord hemisection (C2HS) SCI on the right side, anesthetized with isoflurane (Piramal Critical Care, USA; ∼1.5% in O2). Buprenorphine (endo, USA; 0.03 mg/kg), trimethoprim, sulfadoxine (Somerset, USA; 30 mg/kg), medetomidine (Placadine, USA; 0.1 mg/kg) and carprofen (Rimadyl, 5 mg/kg) were administered subcutaneously 10-20 min before inducing isoflurane anesthesia in a closed chamber (5% isoflurane in 100% O2). Rats were orotracheally intubated and ventilated with a rodent ventilator (model 683; Harvard Apparatus, South Natick, MA, USA). Anesthesia was maintained throughout the procedure (1.5–2% isoflurane in 100% O2). Skin and muscles were retracted, and a C2 laminectomy and durotomy were performed. For C2HS animals, the spinal cord was hemisected unilaterally just caudal to the C2 dorsal roots with micro-scissors, followed with a micro-scalpel to ensure complete sectioning of all remaining fibers. Muscle and skin were then sutured. At the conclusion of the procedure, rats were administered antisedan (1.2 mg/kg, s.q.; Zoetis, Parsippany-Troy Hills, New Jersey) to reverse medetomidine, and lactated Ringer’s solution (4 ml, s.q.) was also provided. Isoflurane was discontinued, the endotracheal tube was removed, and the rats were monitored throughout recovery. A representative transverse image of C2 hemisected spinal cord is shown in Figure 1E.

**Figure 1:**
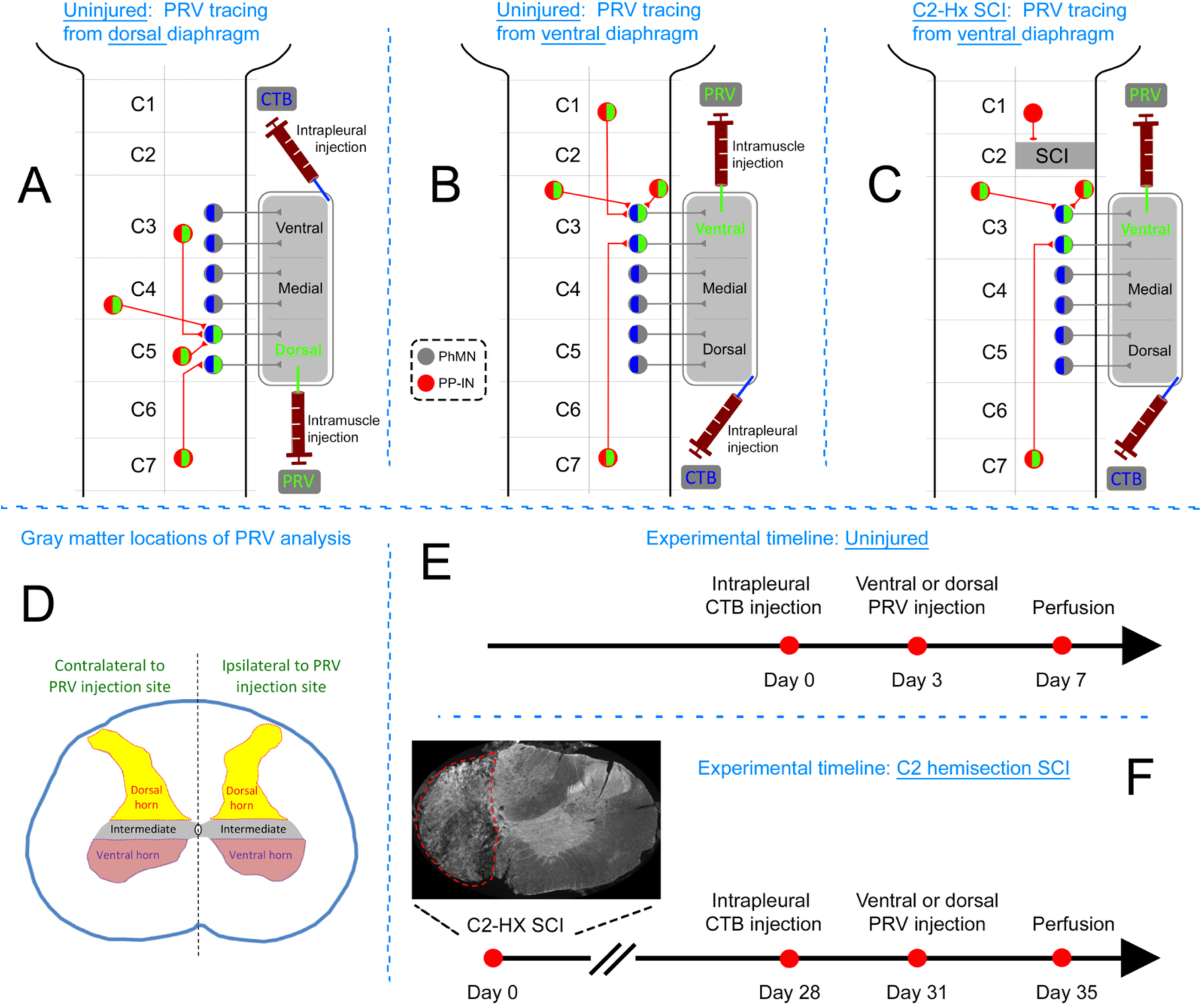
Experimental design. Schematics of three experimental groups: uninjured rats with PRV injection selectively into dorsal diaphragm (**A**); uninjured rats with PRV injection selectively into ventral diaphragm (**B**); C2HS rats uninjured rats with PRV injection selectively into ventral diaphragm (**C**); rats in all groups received intrapleural CTB injection. At each cervical spinal cord segment, PRV-labeled PP-INs were quantified in dorsal horn, intermediate gray matter and ventral horn, both ipsilateral and contralateral to the site of PRV injection (**D**). Experimental timelines for uninjured (**E**) and C2HS (**F**) animals. Representative transverse image of C2HS site shows anatomical completeness of the C2HS lesion (**F**).

### Retrograde labeling of phrenic motor neurons (PhMNs)

Seven days before euthanasia, all rats were anesthetized with isoflurane in 100% O2 balanced (5% for induction; 2-2.5% for maintenance) and 15–20 μL cholera toxin B subunit (CTB, #104 List Biological Labs, Campbell, California, USA) was bilaterally injected into the intrapleural space to retrogradely label PhMNs, as previously described [23].

### Retrograde trans-synaptic tracing of phrenic motor network with pseudorabies virus (PRV)

Four days before euthanasia, rats were anesthetized with isoflurane in 100% O2 balanced (5% for induction; 2-2.5% for maintenance). Animals were placed on a heating pad to maintain a constant body temperature at 37.5 ± 0.5 °C. The depth of anesthesia was confirmed by the absence of response to toe pinch. A laparotomy was performed, and the liver was gently moved dorsally to access diaphragm. Gauze soaked with warm phosphate-buffered saline was placed on the liver to prevent dehydration. Three µL of PRV-152 (PRV-Bartha, Princeton Neuroscience Institute Viral Core Facility, produced on 01/21/25) were injected intramuscularly into the ventral (n = 19; for both uninjured and C2HS) or dorsal (n = 7) portion of diaphragm. Tissues were sutured back and rats were given lactated Ringer’s solution (4 ml, s. q.) and buprenorphine (0.01 mg/kg, s.q.) (Patterson Veterinary, Greeley, Colorado).

### Tissue processing

Four days after PRV injection, rats were intracardially perfused with heparinized saline (0.9% NaCl), followed by 4% paraformaldehyde. Brain and spinal cord were dissected out and post-fixed at 4°C for 24h in 4% PFA. Tissues were then cryoprotected for 48 h in 30% sucrose (in 1X PBS).

### Immunohistochemistry

Frozen transverse (n = 24) or sagittal (n = 2) spinal cord sections (30 µm) were cut using a Microm HM550 cryostat (Thermo Fisher Scientific, Waltham, MA, USA). Free-floating sections were then stored at −20°C in cryoprotectant solution (sucrose 30%, ethylene glycol 30% and PVP40 1% in PBS 1X). Spinal cord sections were rinsed (3 x 5 min) in 1X PBS, blocked for 30 min in blocking solution (5% normal donkey serum (NDS) and 0.2% triton in 1X PBS), and then incubated overnight at 4°C in primary antibody solution (5% NDS and 0.1% triton in 1X PBS). The primary antibody used was rabbit-anti-dsRed (1:500, Living Colors DsRed pAb, RRID: AB_10013483; Takara Bio USA, Mountain View, CA, USA) to enhance PRV labeling visualization. Sections were washed 3 x 5 min in 1X PBS and then incubated in secondary antibody solution (5% NDS and 0.1% triton in 1X PBS) for 2h at room temperature. The secondary antibody used was donkey anti-rabbit Alexa Fluor 647 (1:2000, A31573 Invitrogen, Carlsbad, CA, USA). Sections were washed 3 x 5 min in 1X PBS, mounted on slides, and coverslipped using mounting medium. PRV-labeled PP-INs were counted using a Leica epifluorescence microscope (Axio Imager M2), and representative images were captured with a Leica SP8 confocal microscope (Leica Microsystems Inc., Buffalo Grove, IL, USA) and LAS X imaging software (RRID: SCR_013673).

### Data processing and statistical analyses

All data were presented as mean ± SD. Statistics were considered significant when p < 0.05. RStudio 2026.06.0+242 was used for statistical analyses, and GraphPad Prism 11.0.2 was used for graphing. Cell counts were analyzed in R using separate generalized linear mixed-effects models for each anatomical region. Percentages for heatmaps were analyzed in R using PERMANOVA analyses (ipsilateral, contralateral and central area were analyzed together).

### Data availability

All raw data from this study will be deposited to the Open Data Commons for Spinal Cord Injury (ODC-SCI) for open sharing of data according to the FAIR data principles.

## RESULTS

### PRV tracing from diaphragm revealed the localization of PP-INs throughout cervical spinal cord

Given the extensive diversity of spinal interneurons in type, localization and function within spinal cord [10], we performed neuroanatomical analysis of intrasegmental and propriospinal PP-IN circuitry, including in the context of the topographic innervation of diaphragm by PhMNs. While the PhMN pool extends as a continuous column across C3 to C5/6 ventral horn, previous work has shown that the more caudally located PhMNs preferentially innervate the dorsal subregion of diaphragm muscle, while the more rostrally located PhMNs differentially innervate ventral diaphragm. To investigate PP-IN circuitry, we separately traced PP-IN connections to PhMNs that innervate different portions of the diaphragm. Specifically, we unilaterally applied (via targeted intra-muscle microinjection) trans-synaptic PRV-152 tracer selectively to either dorsal (Figure 1A) or ventral (Figure 1B) diaphragm in uninjured rats. In addition, we unilaterally applied PRV to ventral diaphragm in C2HS rats (Figure 1C) to investigate the impact of cervical SCI on plasticity of PP-IN circuitry compared to the intact condition. In all rats, we also performed intrapleural injection of the non-trans-synaptic retrograde tracer, CTB, to label the cell bodies of only PhMNs throughout the phrenic nucleus. We quantified numbers of PRV-labeled PP-INs individually at segments across cervical spinal cord, including both ipsilateral and contralateral to the side of PRV delivery. Furthermore, we performed this analysis separately in dorsal horn, intermediate gray, and ventral horn at each cervical spinal cord level (Figure 1D).

In sagittal sections from uninjured rats, PRV-labeled PhMNs were found primarily in the rostral C3-C4 portion of the phrenic nucleus when PRV was selectively injected into ventral diaphragm (Figure 2A, D), while PRV-labeled PhMNs were located mostly in caudal C5-C6 portion of the PhMN pool when PRV was instead delivered to dorsal diaphragm (Figure 2B, E). As shown in transverse sections, there was significant co-labeling of PRV and CTB in ventral horn within C3-C6 spinal cord (Figure 2C, F-K), demonstrating that PhMNs were efficiently infected with PRV, allowing for subsequent trans-synaptic transfer to pre-synaptic PP-INs.

**Figure 2:**
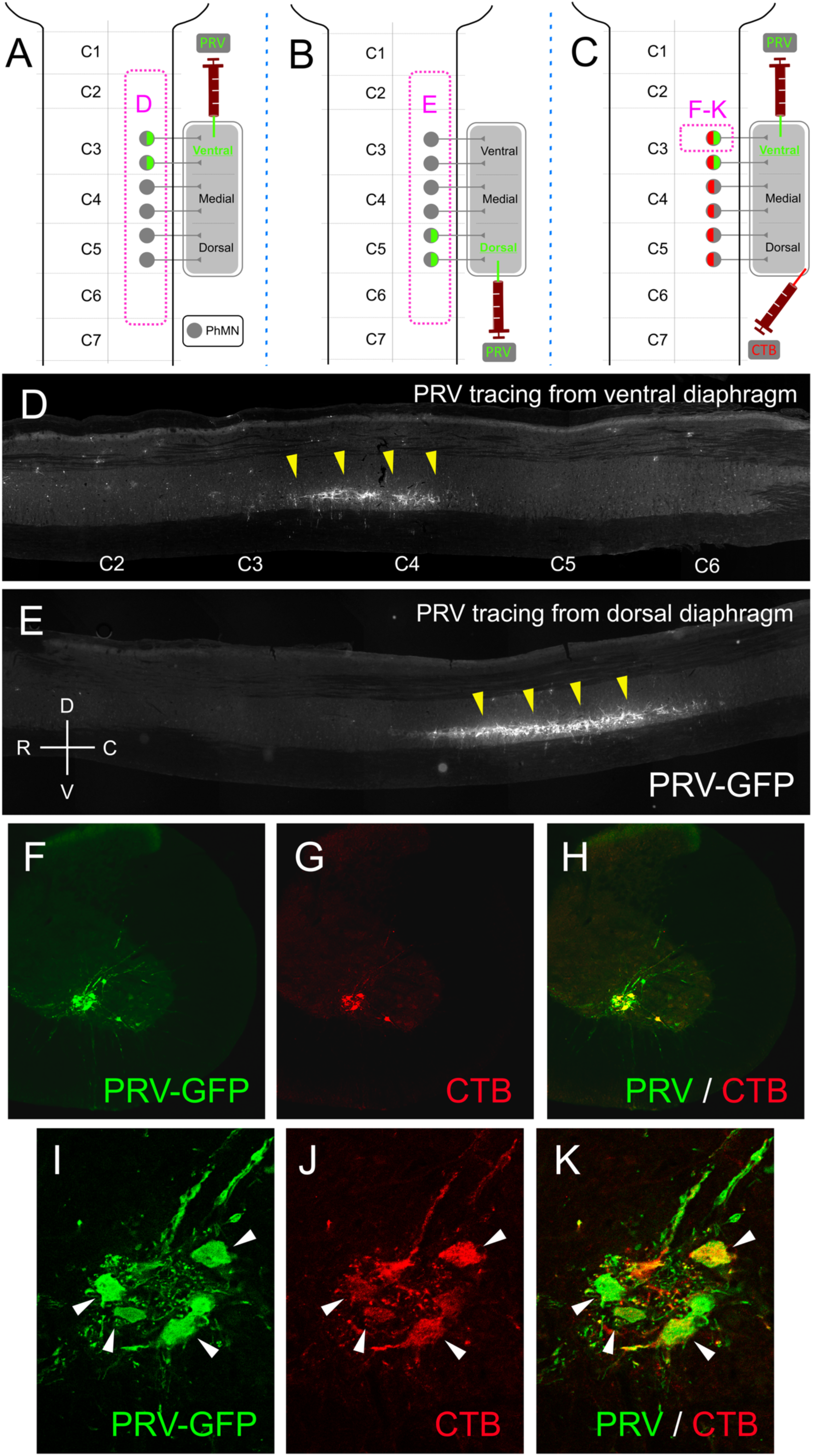
PRV and CTB labeling of PhMNs. (**A**) Schematic showing location of imaging in panel **D**. (**B**) Schematic showing location of imaging in panel **E**. (**C**) Schematic showing location of imaging in panels **F**-**K**. Sagittal section showing PRV-labeled PhMNs following unilateral PRV injection into ventral diaphragm of uninjured animal (**D**). Sagittal section showing PRV-labeled PhMNs following unilateral PRV injection into dorsal diaphragm of uninjured animal (**E**). Directions in panel **E**: D = dorsal; V = ventral; R = rostral; C = caudal. Yellow arrowheads in **D**-**E** denote PRV-labeled PhMNs. Transverse section showing PhMNs in ventral horn of uninjured animal co-labeled with PRV and CTB (**F**-**H**). Higher magnification of transverse image (**I**-**K**); arrowheads denote co-labeled cells.

In uninjured rats, we observed PRV-labeled PP-INs localized widely across cervical spinal cord segments ranging from C1 to C7, including both ipsilateral and contralateral to PRV injection. We also found these PP-INs in various grey matter locations, including dorsal horn, intermediate gray, ventral horn, and surrounding the central canal. Representative PRV-labeled PP-INs from uninjured animals that received PRV injection into ventral diaphragm are shown in Figure 3. Large numbers of at-level projecting intrasegmental PP-INs were found across gray matter laminae of C3 spinal cord following ventral diaphragm PRV injection (Figure 3A-D). Interestingly, we also observed PP-INs that physically interact with the central canal through a process/protrusion (Figure 3A, E-F), with most of these cells found at C3-C6. In addition to intrasegmental PP-INs, we also found significant numbers intersegmentally projecting propriospinal PP-INs. As an example, in uninjured rats that received ventral diaphragm injection of PRV, PP-INs were located at levels throughout cervical spinal cord, including rostrally at C1 and C2 (Figure 3G-J). In addition, while PRV-labeled PP-Ins were mostly separated from one another, we also occasionally observed clusters of PP-INs (Figure 3I). Interestingly, these clusters were only seen in the ipsilateral intermediate gray matter at C1 and C2 levels.

**Figure 3:**
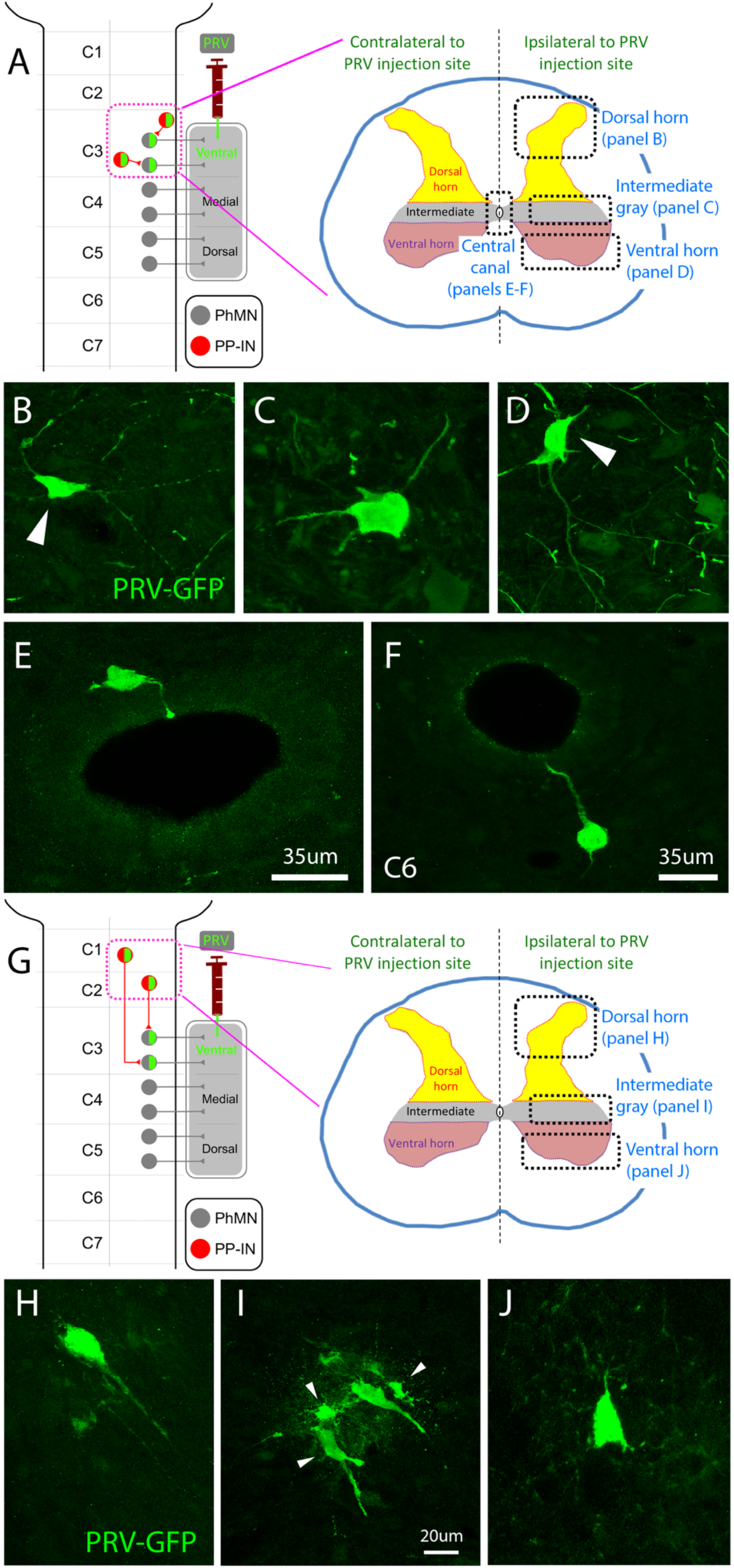
Trans-synaptic tracing of PP-INs in cervical spinal cord. (**A**) Schematic showing location of imaging in panels **B**-**F**. Uninjured rats were injected with PRV injection selectively into ventral diaphragm, and intrasegmental projecting PP-INs were imaged at C3. PRV-labeled PP-INs in C3 spinal cord ipsilateral to PRV injection site in dorsal horn (**B**), intermediate gray matter (**C**), and ventral horn (**D**), as well as directly surrounding the central canal (**E**-**F**). (**G**) Schematic showing location of imaging in panels **H**-**J**. Uninjured rats were injected with PRV injection selectively into ventral diaphragm, and propriospinal PP-INs were imaged at C1/C2. PRV-labeled PP-INs in C1/C2 spinal cord ipsilateral to PRV injection site in dorsal horn (**H**), intermediate gray matter (**I**), and ventral horn (**J**).

### Ventral versus dorsal diaphragm PRV injection showed differential localization of PP-INs

By comparing PRV injection into dorsal versus ventral diaphragm of uninjured animals, we aimed to determine whether PP-IN distribution within cervical spinal cord varied depending on the subpopulation of PhMNs (i.e., rostral-caudal position of the PhMN within the phrenic nucleus and the subregion of diaphragm the PhMN innervates) to which they are synaptically connected. Quantification of PRV-labeled PP-IN localization is presented in heatmaps to illustrate PP-IN distribution across cervical spinal cord segments, across gray matter laminae, and in spinal cord both ipsilateral and contralateral to the PRV injection site for all three experimental groups.

For both dorsal and ventral diaphragm PRV injection into uninjured rats, while some PP-INs were located in the contralateral cervical spinal cord (Figure 5), the vast majority were instead present ipsilateral to the side of PRV delivery (Figure 4). Following dorsal diaphragm PRV injection, labeled PP-INs were found at the highest numbers in various gray matter locations in C5-C6 (Figure 4A). On the contrary, animals with ventral diaphragm PRV injection showed PP-INs localized mostly in C3-C4 (Figure 4B). We also found similar results for the rostral-caudal distribution of PP-INs directly surrounding the central canal following dorsal (Figure 4D) versus ventral (Figure 4E) diaphragm PRV injection. These findings demonstrate that a large portion of PP-INs is intrasegmentally connected to PhMNs and also show that the position of PP-INs closely matches the rostral-caudal location of their connected PhMNs within the phrenic nucleus.

**Figure 4:**
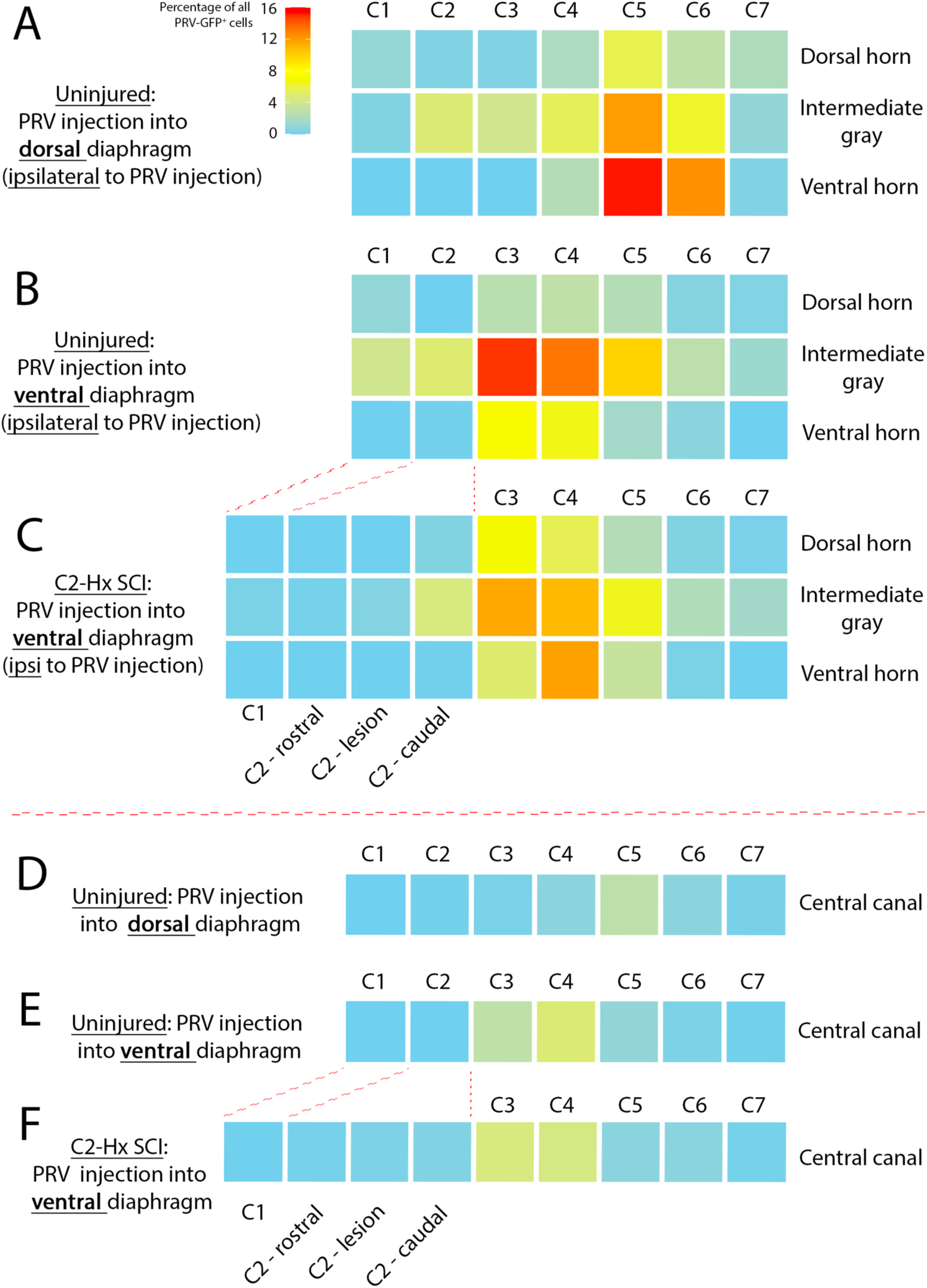
Heatmaps showing distribution of PRV-labeled PP-INs in cervical spinal cord of uninjured and C2HS rats on ipsilateral side and in central canal area. Heatmaps displaying distribution of PRV-labeled PP-INs across the ipsilateral C1-C7 spinal cord: uninjured with dorsal diaphragm PRV injection **(A)**; uninjured with ventral diaphragm PRV injection **(B)**; C2HS with ventral diaphragm PRV injection **(C)**. Heatmaps displaying distribution of PRV-labeled PP-INs directly surrounding the central canal across the C1-C7 spinal cord : uninjured with dorsal diaphragm PRV injection **(D)**; uninjured with ventral diaphragm PRV injection **(E)**; C2HS with ventral diaphragm PRV injection **(F)**. Statistics were performed using PERMANOVA analysis: Dorsal vs ventral injection: p = 0.0024. * p = 0.045. Uninjured vs C2HS: p = 0.3662.

**Figure 5:**
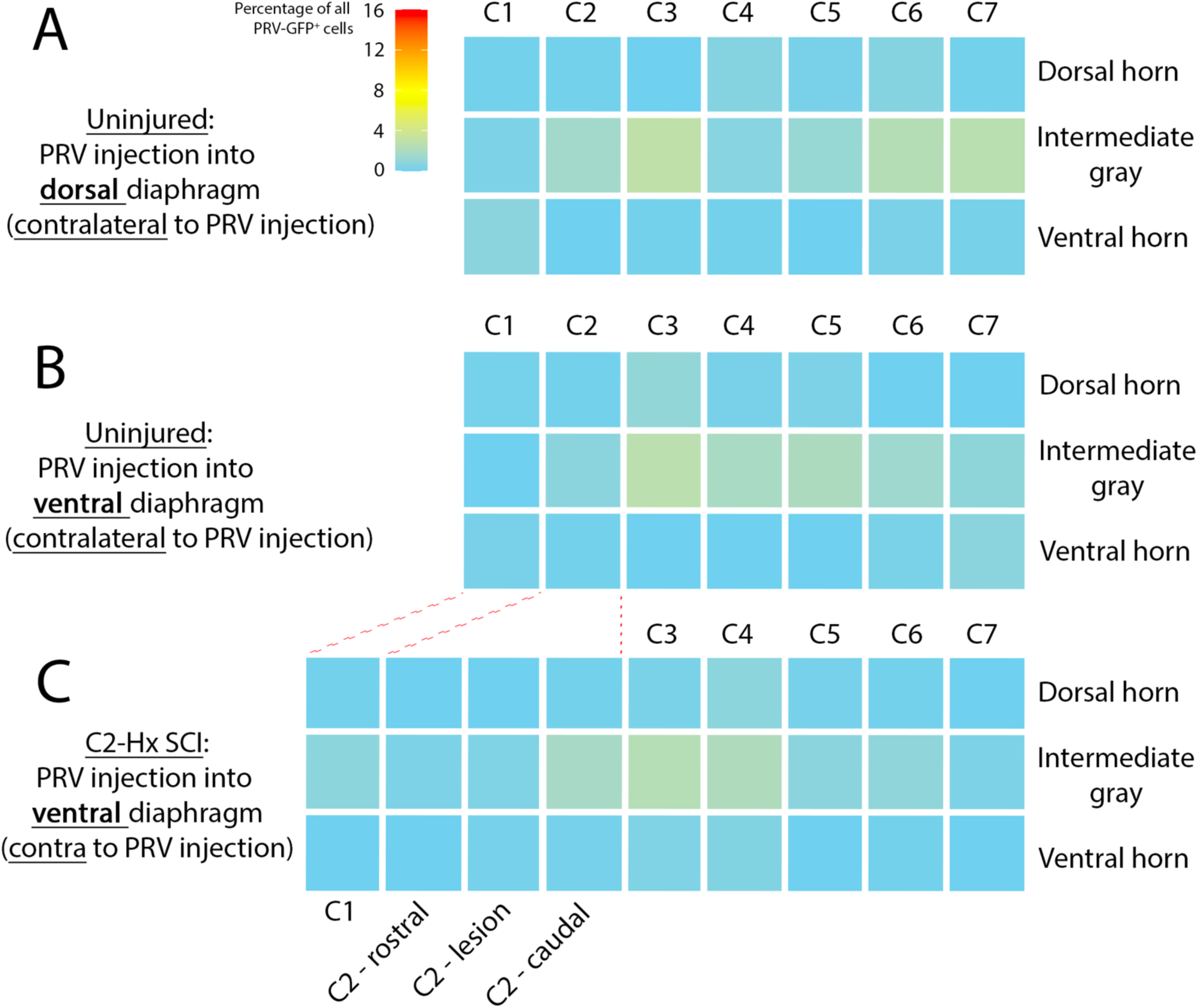
Heatmaps showing distribution of PRV-labeled PP-INs in cervical spinal cord of uninjured and C2HS rats on contralateral side. Heatmaps displaying distribution of PRV-labeled PP-INs across contralateral C1-C7 spinal cord: uninjured with dorsal diaphragm PRV injection **(A)**; uninjured with ventral the diaphragm PRV injection (**B**); C2HS with ventral diaphragm PRV injection (**C**). Statistics were performed using PERMANOVA analysis: Dorsal vs ventral injection: p = 0.0024. * p = 0.0024. Uninjured vs C2HS: p = 0.3662.

We also found differential localization within gray matter laminae depending on the site of PRV injection. PP-INs were more concentrated in C5-C6 ventral horn after dorsal diaphragm PRV injection (Figure 4A), while they were more concentrated in C3-C4 intermediate grey matter with PRV injection into ventral diaphragm (Figure 4B).

Though a greater portion of PP-INs appears to be intrasegmentally projecting, we also observed significant numbers of propriospinal PP-INs with PRV injection into both diaphragm regions. Similar to the differential rostral-caudal distribution of at-level projecting PP-INs, we also found differences in location of these propriospinal PP-INs. Specifically, in animals with dorsal diaphragm PRV injection, PP-INs were located intersegmentally at C2-C4 (Figure 4A), while ventral PRV injection resulted in PP-IN identification with a more rostral distribution at C2 and significantly also at C1 (Figure 4B). Furthermore, it appeared that the vast majority of propriospinal PP-INs were located in intermediate gray matter in both PRV injection conditions, with very few observed in dorsal horn, ventral horn or directly surrounding the central canal.

Together, these findings highlight the differential rostral-caudal and dorsal-ventral organization of PP-INs within the intact cervical spinal cord, including how this PP-IN positioning relates to location of the PhMN subpopulation to which a PP-IN is synaptically connected.

### Cervical hemisection SCI resulted in limited alterations in intrasegmental PP-IN connectivity, but significantly damaged connectivity of propriospinal PP-INs with PhMNs

To investigate plasticity of PP-IN circuitry in response to cervical SCI, rats receiving C2HS were injected with PRV selectively into ipsilesional ventral diaphragm, and these animals were compared to uninjured rats that similarly received ventral diaphragm PRV delivery. C2HS results in significant disruption in synaptic input to PhMNs from neuron populations located rostral to the lesion [16]. This is reflected in our data, in particular with respect to connectivity of propriospinal PP-IN populations. As described above, propriospinal PP-INs were located primarily in intermediate gray matter of C1 and C2 in uninjured animals with ventral diaphragm PRV injection. In C2HS rats, PRV labeling of these propriospinal PP-INs was fully lost in both C1 and in the portion of C2 rostral to the lesion site (Figure 4C), demonstrating that C2HS completely synaptically disconnects propriospinal PP-INs located at high cervical levels with ipsilesional PhMNs.

In general, other than disconnection of PhMNs from C1-C2 propriospinal PP-INs, the PRV-labeled PP-IN distribution pattern in ipsilateral cervical spinal cord between uninjured and C2HS rats did not significantly vary, though there were some minor alterations in localization at various locations (Figure 4B versus 4C; Figure 4E versus 4F). In addition, following C2HS, there were still very few PRV-labeled PP-INs in contralesional spinal cord (Figure 5C), similar to the uninjured condition.

In addition to the heatmaps, quantification of PRV-labeled PP-IN localization in ipsilateral (Figure 6) and contralateral (Figure 7) cervical spinal cord is presented in bars graphs that include individual data points for each animal that received ventral diaphragm PRV injection. Statistical analysis did not highlight differences between uninjured and C2HS conditions, though statistical differences in the number of PP-INs were found across spinal cord segments within an experimental group, as discussed above and as illustrated by the heatmaps.

**Figure 6:**
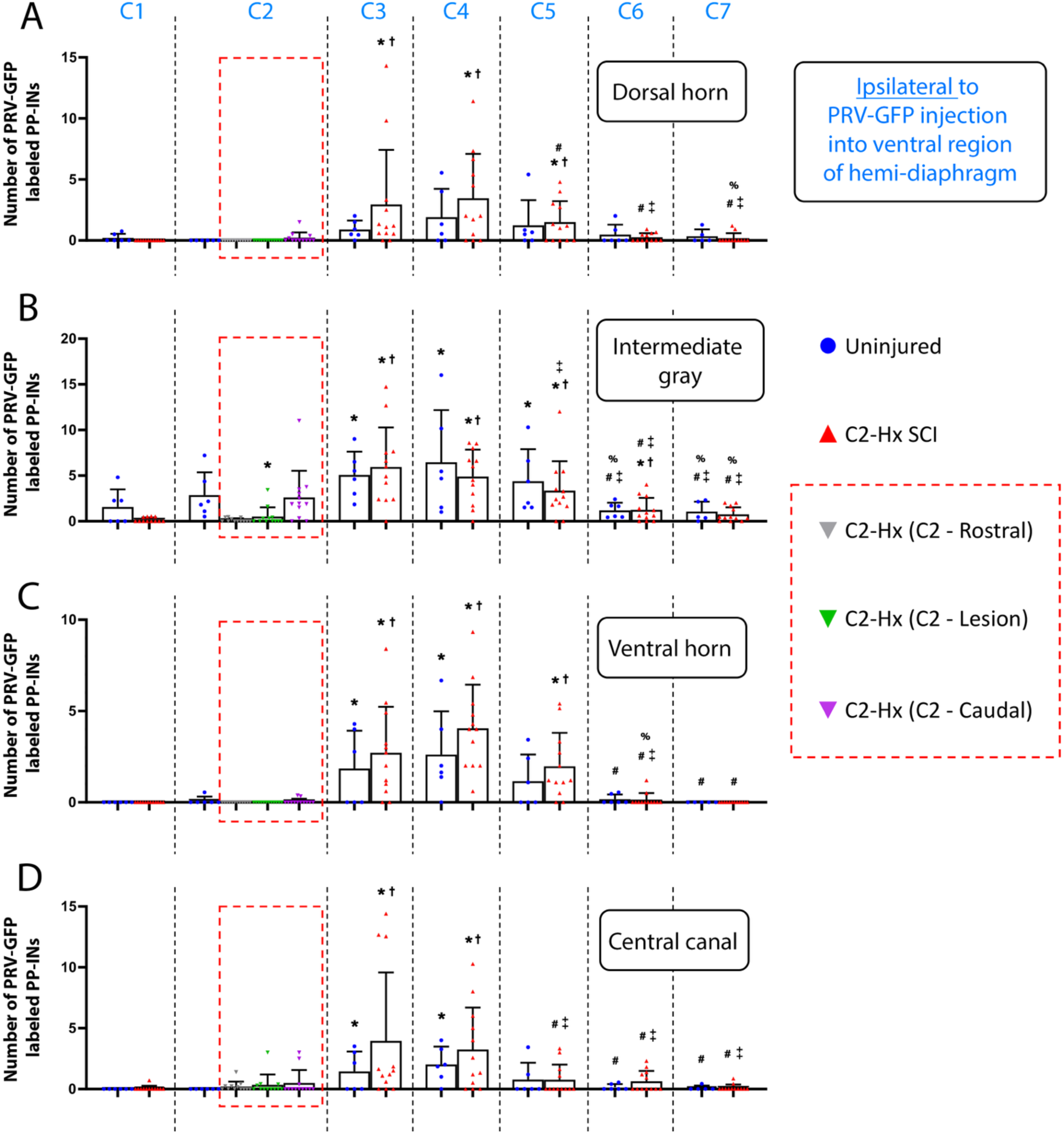
Quantification showing distribution of PRV-labeled PP-INs in cervical spinal cord of uninjured and C2HS rats on ipsilateral side and in central canal area. Bar graphs showing distribution of PRV-labeled PP-INs across the ipsilateral C1-C7 spinal cord and directly surrounding the central canal in both uninjured rats and C2HS rats with ventral diaphragm PRV injection: dorsal horn (**A**); intermediate gray matter (**B**); ventral horn (**C**); directly surrounding central canal (**D**). C2HS - dorsal horn: * p < 0.05 versus C1, † p < 0.05 versus C2, ‡ p < 0.05 versus C3, # p < 0.05 versus C4, % p < 0.05 versus C5. Uninjured - intermediate grey : * p < 0.05 versus C1, ‡ p < 0.05 versus C3, # p < 0.05 versus C4, % p < 0.05 versus C5. C2HS - intermediate grey : * p < 0.05 versus C1, † p < 0.05 versus C2, ‡ p < 0.05 versus C3, # p < 0.05 versus C4, % p < 0.05 versus C5. Uninjured - ventral horn: * p < 0.05 versus C1, # p < 0.05 versus C4. C2HS - ventral horn: * p < 0.05 versus C1, † p < 0.05 versus C2, ‡ p < 0.05 versus C3, # p < 0.05 versus C4, % p < 0.05 versus C5. Uninjured - central canal area: * p < 0.05 versus C1, # p < 0.05 versus C4. C2HS – central canal area: * p < 0.05 versus C1, † p < 0.05 versus C2, ‡ p < 0.05 versus C3, # p < 0.05 versus C4. Statistics were performed using a binomial generalized linear mixed model (GLMM). No statistical differences were observed for uninjured vs C2HS: p > 0.05.

**Figure 7:**
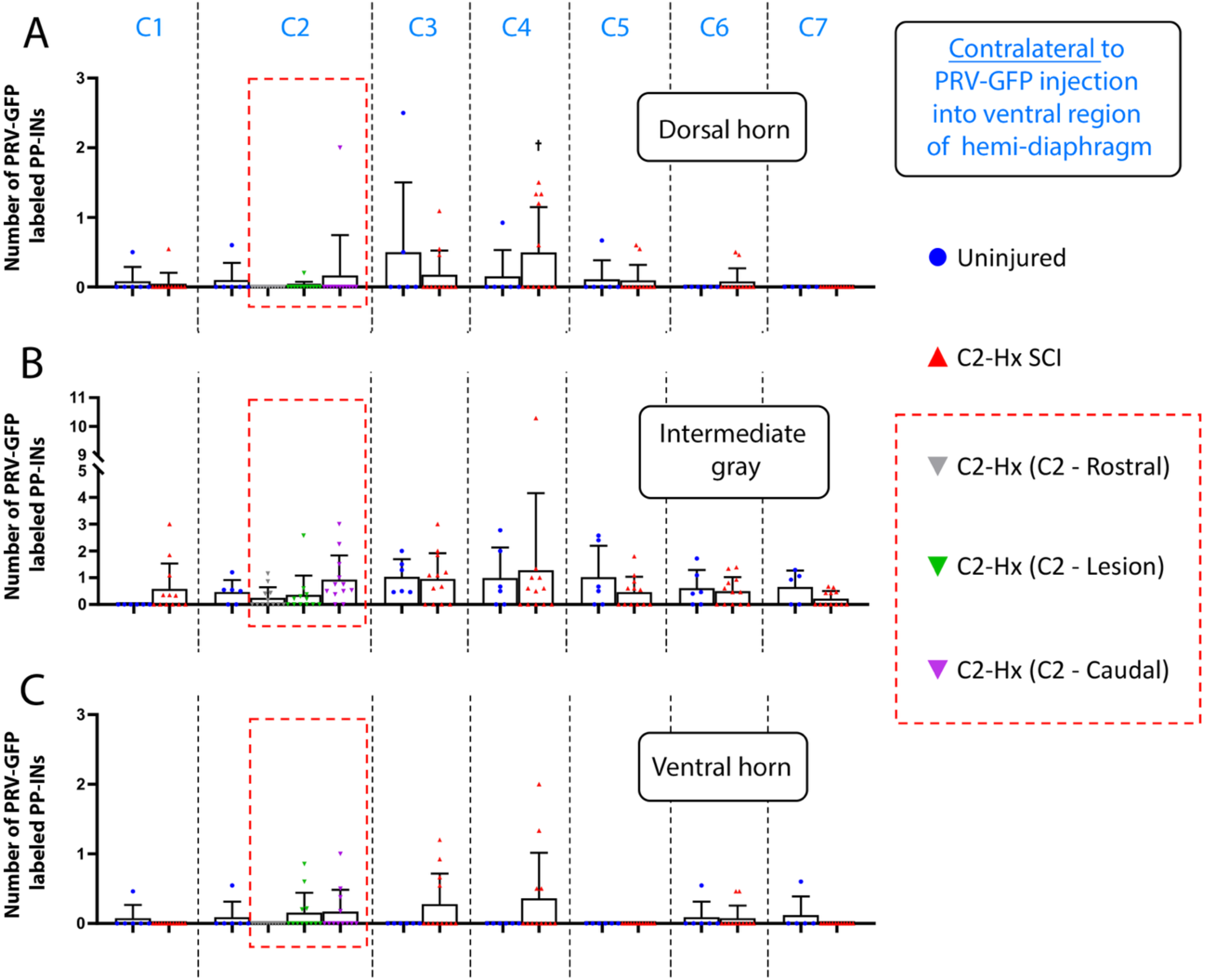
Quantification showing distribution of PRV-labeled PP-INs in cervical spinal cord of uninjured and C2HS rats on contralateral side. Bar graphs showing distribution of PRV-labeled PP-INs across the contralateral C1-C7 spinal cord in both uninjured rats and C2HS rats with ventral diaphragm PRV injection: dorsal horn **(A)**; intermediate gray matter **(B)**; ventral horn **(C)**. C2HS - dorsal horn: † p < 0.05 versus C2. Statistics were performed using a binomial generalized linear mixed model (GLMM). No statistical differences were observed for uninjured vs C2HS: p > 0.05.

Taken together, these findings demonstrate that C2HS results in limited alterations in the connectivity of intrasegmental PP-IN circuity, while propriospinal PP-IN to PhMN circuitry is dramatically damaged.

## DISCUSSION

This work was aimed at providing enhanced understanding of PP-IN circuitry within the intact cervical spinal cord, as well as how this circuitry is impacted by cervical SCI. We observed that a portion of the PRV-labeled cells within cervical spinal cord were PhMNs (co-labeled with CTB) residing within C3-C6 ventral horn. This was expected since PhMNs innervate diaphragm and therefore serve as the PRV entry point into spinal cord. The PRV injection paradigm (ventral versus dorsal diaphragm injection of PRV) allowed us to observe differential labeling of PhMNs: primarily at C3-C4 following ventral diaphragm PRV injection and mostly at C4-C6 for dorsal injection. This observation is in line with previous work in cat by Laskowski and Sanes who showed that the ventral subregion of diaphragm muscle is primarily innervated by PhMNs located at C3, medial subregion is innervated by C4 PhMNs, and dorsal subregion is controlled by PhMNs located at the C5 level [24]. However, the question about PP-IN localization in the context of this topographic innervation of diaphragm by PhMNs remained unanswered. Here, we addressed several points. With both ventral and dorsal diaphragm injection, we observed PRV-labeled PP-INs throughout C1 to C7; therefore, regardless of where PhMNs are located, they can connect to PP-INs throughout the entire extent of cervical spinal cord. A study by the Lane group reported a similar distribution of premotor INs connected to forelimb motor neurons. This finding suggests that PP-INs may follow similar organizational pattern as other pre-motor INs populations: they reside across multiple spinal cord segments, are located both ipsilaterally and contralaterally (to the initial PRV injection site), and are found throughout the spinal grey matter [25].

As the aim of this study was to characterize anatomical distribution of PP-INs within cervical spinal cord, determining their phenotype and specific role within the phrenic network was beyond the scope of this work and is beginning to be addressed by our and other groups [10, 14]. A number of studies have described a V2a subpopulation of excitatory PP-INs, though inhibitory and excitatory PP-INs co-exist within the phrenic neuronal network [20, 26–28]. The Crone group showed these V2a PP-INs (located within brainstem and cervical spinal cord) in adult mice are involved in organizing and distributing respiratory motor output across different respiratory muscles but do not seem to provide essential tonic motor inspiratory drive during eupnea [27]. The potential role of other types of PP-INs in eupneic breathing remains to be investigated. Recent work by the Lane and Zholudeva groups also suggests V2a PP-INs can be recruited as a form of respiratory plasticity following high SCI [20]. Building on these findings, they targeted these PP-INs as a therapeutic tool by transplanting neural progenitor cells enriched with V2a INs into injured C3-C4 spinal cord. Transplanted cells survived and integrated into host phrenic circuitry and were associated with partial recovery of diaphragm activity [29].

Regarding plasticity of PP-IN connectivity following C2HS, we did not observe significant reorganization, contrary to what the Lane group found for PP-INs [20], but similar to what they observe for lumbar premotor INs in the same C2HS model [25]. For PP-INs, they showed that the number of labeled PP-INs was greater within the contralesional (but not ipsilesional) side two weeks following C2HS compared to uninjured rats. Specific identification of connected Chx10 neurons (V2a PP-INs) revealed their numbers are significantly increased in C2HS rats on ipsilesional and contralesional sides [20]. The difference observed with our data may be due to the delayed post-injury and post-PRV injection temporal design we employed; we performed evaluation at 35 days post-injury and 92 hours post-PRV injection, while the Lane group performed their study at 14 days post-injury and 72 hours post-PRV injection. In addition, we counted PP-INs separately at each spinal cord segment, while they conducted PP-IN quantification across all segments.

We also made some interesting additional observations. While most observed PP-INs were positioned individually, some seemed to be organized in clusters, specifically at C1-C2 levels. These were consistently observed in intermediate grey matter and only unilaterally, suggesting the possible existence of a conserved upper cervical microdomain. Though this region overlaps the central cervical nucleus [30] and lies near the reported distribution of upper cervical inspiratory neurons [31], their identity and functional significance remains to be determined.

The observed labeling of PP-INs far caudally at C7 (including with PRV injection into both dorsal and ventral diaphragm) suggests the phrenic network receives ascending inputs from (and/or sends descending outputs toward) lower spinal cord segments. Previous work from the Reier group observed co-labeling of C7 interneurons when they injected rats into diaphragm with PRV-152 and into intercostal muscle with PRV-614, showing these interneurons were connected to both PhMNs and intercostal motor neuron pools [13]. Their results support a role for these interneurons in coordination of respiratory muscles activity. These C7 PP-INs could represent an anatomical interface between respiratory motor output and locomotor and postural activity [32–34].

The observation of PP-INs contacting cerebrospinal fluid (CSF) of the central canal corresponds to described CSF-contacting neurons (CSF-cNs) [35]. In this regard, the Reier group found PP-INs close to the central canal [13], but none making contact with CSF. In other motor neuronal networks, it has been shown that CSF-cNs make direct contact with motor neurons and pre-motor neurons [36]. In addition, the frequency of the locomotor central pattern generator can be decreased when CSF-N activity is chemogenetically inhibited. These studies suggest it is possible that similar CSF-cNs could be identified within the phrenic network. What could be the role of CSF-Ns within the phrenic system since inspiratory frequency/rhythm is generated within brainstem? One hypothesis is they are a monitor of respiratory activity. CSF-Ns are mostly GABAergic, they sense pH in CSF, and cervical CSF-Ns are sensitive to AMPA/kainate glutamate receptor regulation [35]. They could apply an inhibitory control onto the phrenic network and be involved in regulatory processes driven by modulation in CSF pH. More work needs to be done to unravel the potential role of these CSF-Ns in phrenic circuitry.

## Conclusion

In this study, we showed that PP-INs are widespread throughout the C1-C7 spinal cord in the uninjured condition (Figure 8A). These PP-INs have a predominant intrasegmental connectivity pattern with PhMNs, though significant numbers of longer distance projecting propriospinal PP-INs also exist both rostral and caudal to the PhMN pool. Furthermore, while PP-INs connect both ipsilaterally and contralaterally with PhMNs, there is a strong bias to ipsilateral connectivity. C2HS induces a major disconnection between PhMNs and ipsilesional propriospinal PP-INs located rostral to the SCI site (Figure 8B), but conversely induces fairly limited plasticity in the connectivity of intrasegmental projecting PP-INs located within the C3-C6 spinal cord. PP-INs are an important component of respiratory neural circuitry both in the intact nervous system and following SCI. These findings provide important information to aid in targeting this cell type for promoting spinal cord repair.

**Figure 8:**
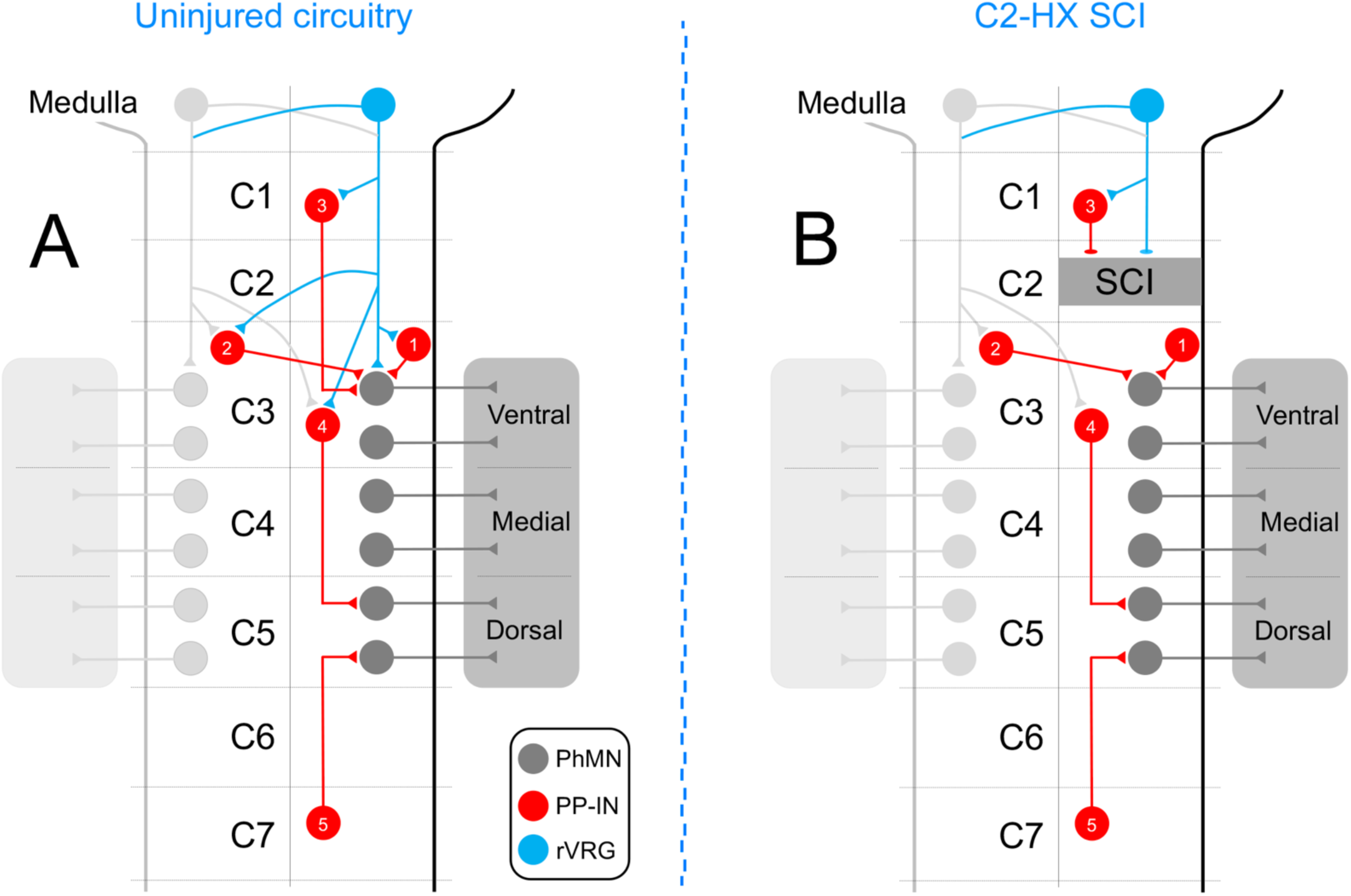
Diagrams summarizing localization of various PP-IN populations in uninjured circuitry and following C2HS. PP-INs are widespread throughout C1-C7 spinal cord in the uninjured condition (**A**). These PP-INs have a predominant intrasegmental projecting connectivity pattern with PhMNs, though significant numbers of longer distance projecting propriospinal PP-INs also exist both rostral and caudal to the PhMN pool. While PP-INs connect both ipsilaterally and contralaterally with PhMNs, there is a strong bias to ipsilateral connectivity. (1) Ipsilateral intrasegmental projecting PP-IN. (2) Contralateral intrasegmental projecting PP-IN. (3) Ipsilateral projecting propriospinal PP-IN with cell body located rostral to the PhMN pool. (4) Ipsilateral projecting propriospinal PP-IN with cell body located within the PhMN pool. (5) Ipsilateral projecting propriospinal PP-IN with cell body located caudal to the PhMN pool. C2HS induces a major disconnection between PhMNs and ipsilesional propriospinal PP-INs located rostral to the SCI site (**B**), but conversely induces fairly limited plasticity in the connectivity of intrasegmental projecting PP-INs located within C3-C6 spinal cord.

## ACKNOWLEDGEMENTS

This work was supported by the National Institute of Neurological Disorders and Stroke (R01NS110385 and R01NS079702 to ACL). We thank the Center for Neuroanatomy with Neurotropic Viruses (CNNV) and the Princeton Neuroscience Institute Viral Core Facility for providing PRV.

## AUTHOR CONTRIBUTIONS (CRediT)

Conceptualization: AC Lepore

Data curation: P Michel-Flutot, AC Lepore

Formal analysis: P Michel-Flutot, AC Lepore

Funding acquisition: AC Lepore

Investigation: P Michel-Flutot, A Harbeck, L Cheng, K Rust, MA Lyttle, SJ Thomas, DA Jaffe, K Crowther, G Daszewska-Smith, GM Smith, S Li, AC Lepore

Project administration: AC Lepore

Supervision: AC Lepore

Validation: P Michel-Flutot, AC Lepore

Visualization: P Michel-Flutot, A Harbeck, L Cheng, K Rust, AC Lepore

Roles/Writing – original draft: P Michel-Flutot, AC Lepore

Writing – review & editing: P Michel-Flutot, AC Lepore

## ABBREVIATIONS

C2, C3, C4, etc.: cervical spinal cord level 2, 3, 4, etc.
C2HS: C2 spinal cord hemisection
SCI: spinal cord injury
PP-IN: pre-phrenic interneuron
PhMN: phrenic motor neuron
PRV: pseudorabies virus
rVRG: rostral ventral respiratory group
CTB: cholera toxin B Subunit
CSF-cNs: cerebrospinal fluid-contacting neurons

